# A behavioral screen for mediators of age-dependent TDP-43 neurodegeneration identifies SF2/SRSF1 among a group of potent suppressors in both neurons and glia

**DOI:** 10.1101/2021.05.12.443894

**Authors:** Jorge Azpurua, Enas Gad El-Karim, Marvel Tranquille, Josh Dubnau

**Affiliations:** Department of Anesthesiology, Stony Brook School of Medicine, NY 11794, USA; Department of Physiology and Biophysics, M.S. Program, Stony Brook School of Medicine, NY 11794, USA; Department of Neurobiology and Behavior, Stony Brook University, NY 11794, USA

## Abstract

Cytoplasmic aggregation of Tar-DNA/RNA binding protein 43 (TDP-43) occurs in 97 percent of amyotrophic lateral sclerosis (ALS), about 40 percent of frontotemporal dementia (FTD) and in many cases of Alzheimer’s disease (AD). Cytoplasmic TDP-43 inclusions are seen in both sporadic and familial forms of these disorders, including those cases that are caused by repeat expansion mutations in the *C9orf72* gene. To identify downstream mediators of TDP-43 toxicity, we expressed human TDP-43 in a subset of *Drosophila* motor neurons. Such expression causes age-dependent deficits in negative geotaxis behavior. Using this behavioral readout of locomotion, we conducted an shRNA suppressor screen and identified 32 transcripts whose knockdown was sufficient to ameliorate the neurological phenotype. The majority of these suppressors also substantially suppressed the negative effects on lifespan seen with glial TDP-43 expression. In addition to identification of a number of genes whose roles in neurodegeneration were not previously known, our screen also yielded genes involved in chromatin regulation and nuclear/import export-pathways that were previously identified in the context of cell based or neurodevelopmental suppressor screens. A notable example is *SF2*, a conserved orthologue of mammalian *SRSF1*, an RNA binding protein with roles in splicing and nuclear export. Our identification *SF2/SRSF1* as a potent suppressor of both neuronal and glial TDP-43 toxicity also provides a convergence with *C9orf72* expansion repeat mediated neurodegeneration, where this gene also acts as a downstream mediator.

**Author Summary:** Loss of nuclear function of TDP-43 and its mislocalization into cytoplasmic inclusions are central features to a suite of neurodegenerative disorders. We screened 2700 *Drosophila* genes to identify downstream mediators that suppress an age-dependent motor dysfunction phenotype when they are knocked down by RNA interference. We identified both previously implicated pathways and several novel genes whose knock down is sufficient to dramatically and robustly rescue TDP-43 toxicity both in neuronal and glial contexts. Notably, we report that *SF2/SRSF1*, which was previously reported as a suppressor of *C9orf72* hexanucleotide expansion repeat toxicity, also is a potent suppressor of TDP-43 mediated neurodegeneration.

## Introduction

Protein aggregation pathology of TDP-43 is a central feature in a suite of neurodegenerative disorders, including ALS, FTD and AD or those with an AD like presentation (Limbic-predominant Age-related TDP-43 Encephalopathy, LATE) [1]. In ALS in particular, TDP-43 protein pathology is seen in ∼97% of patient’s post mortem tissue from affected brain regions [2]. Normally TDP-43 localizes primarily to the nucleus [3] of both neurons and glial cells, where it is plays key roles in splicing, transcriptional regulation, DNA repair and transposable element suppression [4–10]. Thus the pathological mislocalization into cytoplasmic inclusions will involve a loss of nuclear function and may produce toxic cytoplasmic effects as well. Importantly, such pathology is seen across the majority of both familial (fALS) and sporadic (sALS) cases. Familial forms of ALS that exhibit TDP-43 proteinopathy include the rare cases that are caused by mutations in the TDP-43 protein coding region [11,12], and those caused by *C9orf72* repeat expansions [13], which is the most common genetic cause of ALS and FTD in the USA.

To identify mediators of TDP-43 neurodegeneration, several previous studies have capitalized on the toxicity of overexpressing TDP-43. Such over-expression often leads to nuclear clearance and accumulation of cytoplasmic inclusions. This strategy has been successfully employed to identify genes that enhance or suppress TDP-43 aggregation and toxicity, and many of these have been validated in human disease. For instance, screening for suppressors and enhancers in yeast led to the identification of several relevant downstream mediators of TDP-43, FUS and C9ORF72 toxicity [14–17], including ATX2 [18], whose role in human disease epidemiology has since been validated [19,20]. More recently, several suppressor/enhancer screens have been performed in a *Drosophila* developmental context in photoreceptor neurons offering the advantage of being both a metazoan and neuronal context. Together, these studies have identified important core cell biological pathways including dysfunction in nucleocytoplasmic transport [17,21,22] and chromatin remodeling [23] as convergence points of TDP-43 induced pathology.

To identify downstream effectors in the context of an intact neural circuit, we conducted a behavioral screen for suppressors of age dependent decline in negative geotaxis locomotion in *Drosophila*. We used an unbiased RNAi screen of over 2700 genes and identified over 30 that suppressed the locomotion phenotype even in aged animals. This screen identified genetic pathways that have previously been implicated in neurodegeneration, as well as novel robust suppressors not previously connected to TDP-43 toxicity. Among these is the *SF2/SRSF1* gene, which is a known suppressor of C9ORF72 models of ALS/FTD [24].

## Results

### A behavioral screen to identify suppressors of TDP-43-mediated motor neuron toxicity

Expression of a codon-optimized human TDP-43 (*UAS-hTDP-43*) transgene [25] using the E49 motor neuron Gal4 (*E49-Gal4*) driver [26] is sufficient to cause [27] adult fly motor deficits and an increased rate of mortality. We noted that when both the *E49-Gal4* and the *UAS-hTDP-43* were maintained together in a single strain, the motor deficit diminished dramatically over a period of approximately five generations (not shown). This suggested that naturally occurring variants within the genetic background could be selected over time to act as modifiers of TDP-43 toxicity. We therefore postulated that individual gene knock downs might also be used effectively to identify mediators of the toxicity of TDP-43 to motor neurons.

To avoid the confounding impact of selected suppressors emerging among natural variants in the strain, we created a new recombinant stock that contained not only the *E49-Gal4* driver (expression pattern is shown in GFP in **Fig 1A**) and the *UAS-TDP-43* transgene insertion but also a ubiquitously expressed *tub-Gal80* repressor (Gal80) on the X chromosome. The presence of the Gal80 repressor in this strain prevents expression of the *UAS-TDP-43*, and thereby prevented accumulation of modifying alleles within the strain. However, when males of this strain are crossed to wild-type females, the *tub-Gal80* is easily segregated, in male progeny, from the *E49-Gal4* and *UAS-TDP-43* transgenes which reside on autosomes. We found that such male offspring consistently and stably produced severe adult motor defects even after the parental stock was maintained for many (>20) generations. We used this *Gal80;E49>TDP-43* strain to conduct an unbiased genetic screen for suppressors of the adult locomotion defect. By crossing males from this parental strain with females that contained individual UAS-driven shRNAs typically targeting a single gene, we screened the impact of each shRNA on locomotion in approximately one week old male progeny that had lost the Gal80 repressor (**Fig 1B**).

**Fig 1.**
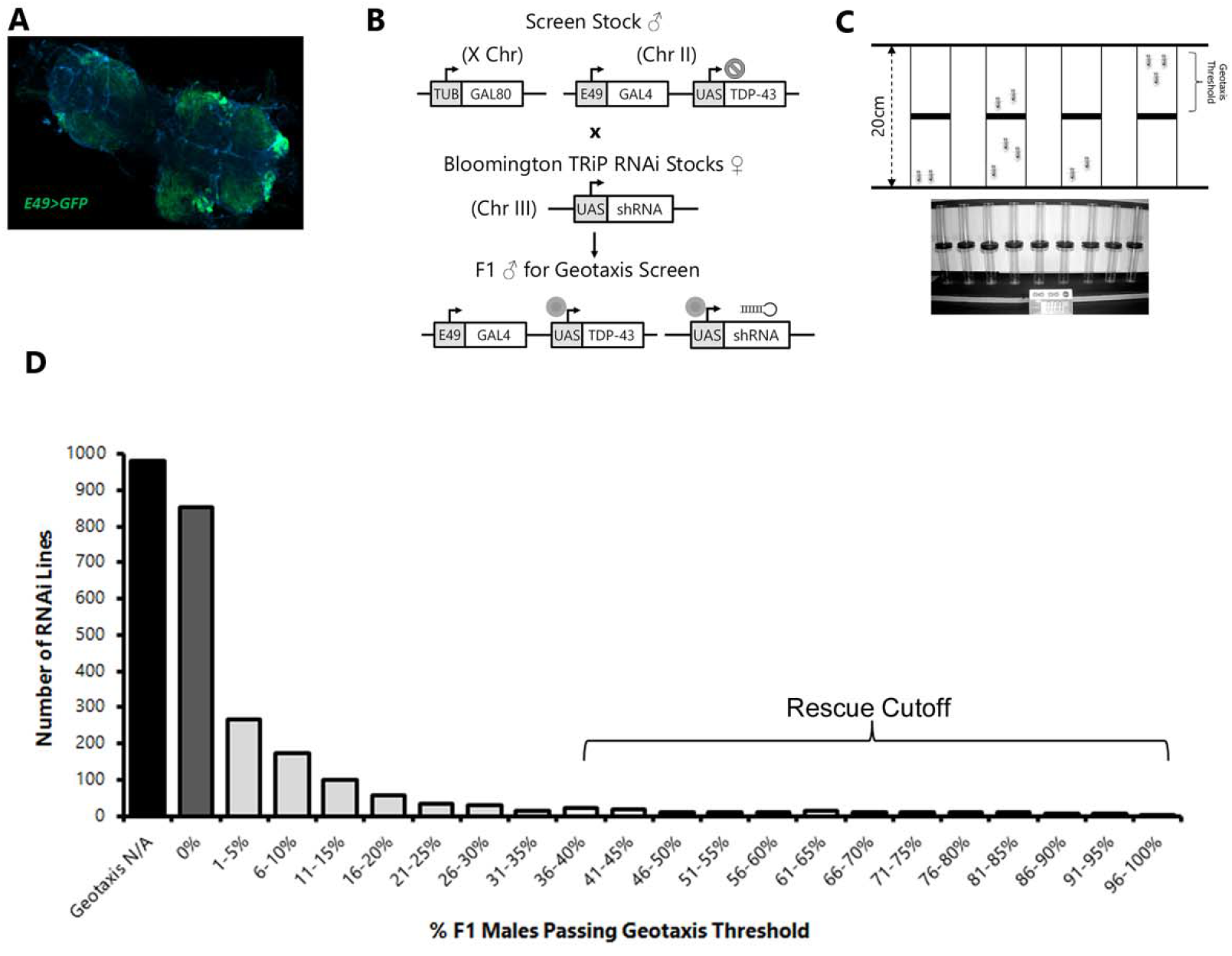
shRNA screen for suppressors of motor-neuron driven TDP-43 negative geotaxis defects. (a) Confocal microscopy image (10x objective) of dissected thoracic ganglion showing GFP expression driven by E49-Gal4. (b) Diagram of cross scheme used in screen. Males from parental strain carrying an X-linked Gal80 and autosomal E49 motor GAL4 driver (subset of motor and sensory neurons) and a codon-optimized human TDP-43 transgene on chromosome II were crossed to virgin females from candidate shRNA lines yielding F1 males lacking GAL80 and therefore expressing TDP-43 and the candidate shRNA in E49 neurons. (c) Diagram and photograph of negative geotaxis setup for behavioral rescue assay. (c) Histogram showing shRNA lines (y-axis) binned by geotaxis performance (% of flies crossing 10cm threshold in 10 seconds, x-axis) across all tested shRNA lines. The line indicates the cutoff for retesting as a candidate suppressor.

The design of this screen involves expressing both the *UAS-TDP-43* and the *UAS-shRNAs* via the same *E49-Gal4* driver. We were cognizant of the fact that the presence of two loci that contain the same Gal4 responsive UAS enhancer might titrate the available Gal4 transcription factor available for each. To mitigate against the possibility that differences in such titration across shRNAs could yield false positive or negative effects, we limited this screen to shRNAs from a single collection that all use an identical “VALIUM20” vector as their RNAi expression system. Additionally, we only used the subset of strains [28] that contained a VALIUM20 *UAS-shRNA* inserted into a specific AttP2 landing site on the third chromosome. This collection of VALIUM20 shRNA insertions at this defined chromosomal location consisted of 2724 strains, generally targeting a specific *Drosophila* genomic locus. We crossed females from each of these 2724 *UAS-shRNA* containing strains with males from our *tub-GAL80;E49>TDP-43* strain (**Fig 1B**). We then quantified negative geotaxis locomotion behavior [29] among F1 male offspring from each cross at approximately 1 week of adult age. In order to scale up the behavioral screening, we designed a simple apparatus (**Fig 1C**) that allowed multiple strains to be tested in parallel. We established a stringent locomotion value (35% of animals crossing 10cm marker within 10seconds) to identify candidate suppressors, and then each candidate suppressor was re-screened with an independent cross to validate the findings from the primary screen (**Fig 1**). This also permitted us to calculate both the overall false positive and false negative rates (**S1 Table, false positive rate ∼2%, false negative rate negligible**). The vast majority of the crosses yielded F1 progeny that had severe locomotion defects. In many cases there either were too few surviving male progeny to screen, or we found that none of the flies were able to climb past the 10cm marker within the allotted time (**Fig 1D; S2 Table**). We identified 33 shRNA crosses (1.24% of total) that resulted in greatly improved locomotion, with greater than 35% of the animals crossing threshold in both the primary screen and the independent validation crosses (**Table 1; S2 Table**). As a further validation, 22 of these screen hits were retested after being equilibrated (**Fig S1**) with 5 generations of back-crossing into the w^1118^(isoCJ1) genetic background (a derivative of Canton S) both to match that of the *GAL80;E49>TDP-43* strain and to test for robustness of the effects in a new genetic background. The suppression of the negative geotaxis behavioral defect persisted with 21/22 tested in this second genetic background (**Table 1**).

**Table 1.**
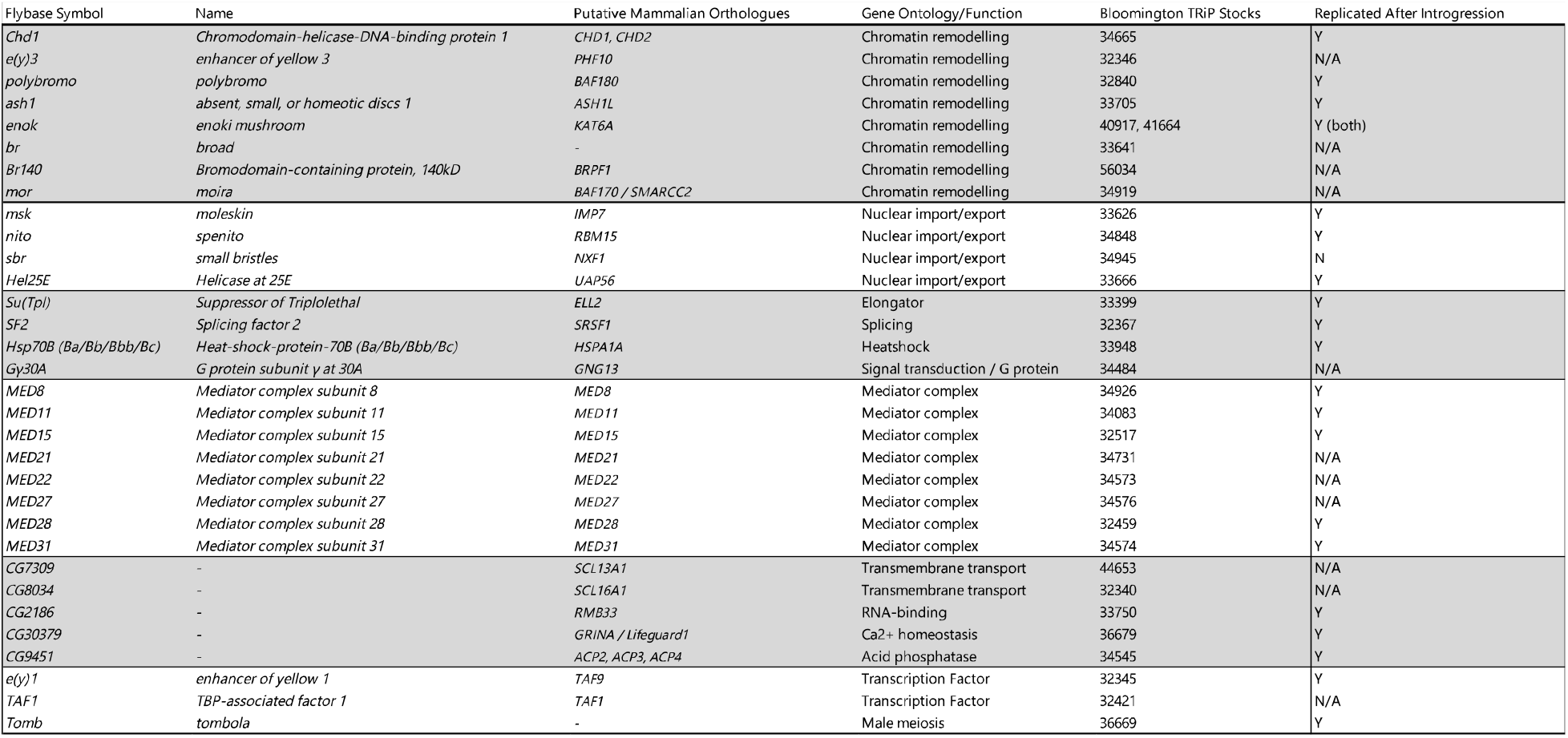

These TDP-43 suppressors fall into several broad functional categories. First, we identified a number of novel genes that have not previously been implicated in neurodegeneration and a relatively large group that are components of the Mediator complex. Mediator proteins integrate signals between transcription factors (TFs) and RNA polymerase, with different subunits showing some specificity for interactions with distinct subsets of TFs [30]. Second are those involved in cell biological functions that are well established to play a role in neurodegeneration. These include chromatin remodeling factors [23,31,32] and those involved in nucleocytoplasmic transport [17,23,33–36]. In the latter category, the msk/importin-7 gene is of interest because the sign of its effect is opposite from what one would predict if it were involved in transport of TDP-43 (see discussion). Finally, we highlight that the SF2/SRSF1 gene was identified among our TDP-43 suppressors. This is notable because this gene has been found as a downstream effector in the context of C9orf72 repeat expansion toxicity [24].

### Motor neuron-specific CRISPR/Cas9 knockout of msk/importin-7 and SF2/SRSF1 suppresses negative geotaxis behavior

We used CRISPR/Cas9 gene editing as an independent means to validate that TDP-43 toxicity to motor neurons can be suppressed by knockout *of SF2/SRSF1* and *msk/importin-7*. (**Fig 2A)** We designed guide RNAs (gRNAs) targeting these genes and we generated transgenic animals that constitutively express these gRNAs by integrating into the same AttP2 landing site that was used for the shRNA screen. As controls, we used gRNAs targeting either GFP or the *lst* gene, which was not identified as a suppressor in our shRNA screen and is not known to be expressed in motor neurons. We tested the impact of expressing each of these gRNAs in animals that also contained the *E49-Gal4* line, *UAS-TDP-43* and *UAS-Cas9* transgenes.

**Fig 2.**
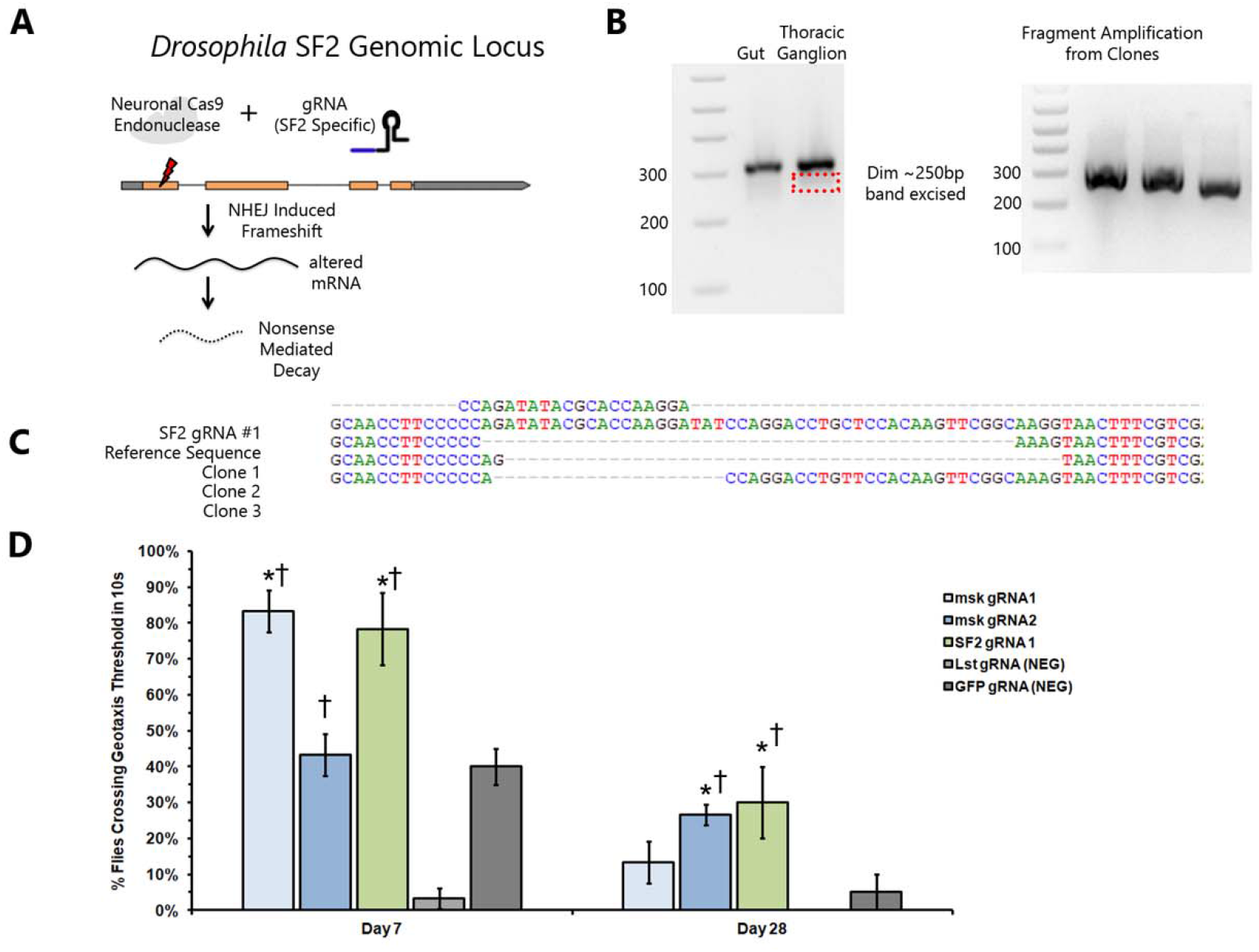
Motor neuron specific CRISPR/CAS9 knock-out of msk/importin-7 and of SF2/SRSF1 are each sufficient to prevent TDP-43 mediated defects in negative geotaxis. (a) Overview of CRISPR/Cas9 knock out strategy. gRNAs specific to *SF2/SRSF1* or *msk/importin-7* (see **S3 Table**) are expressed constitutively in all cells by a U6 promoter plasmid integrated into the AttP2 landing site. Cas9 expression is restricted to E49 neurons, limiting editing to those cells. (b) Gel electrophoresis of PCR products from dissected tissues of male F1 offspring. Gel 1 Lanes: DNA ladder, Gut amplification product (no editing), thoracic ganglion amplification product (editing in some neurons). A faint band corresponding to reduced size predicted from edited products was excised and cloned into a sequencing vector. Gel 2 Lanes: DNA Ladder, and three independent PCR amplifications from sub-cloned edited inserts. These show variable size corresponding to stochastic NHEJ editing of target gene. (c) DNA sequence of edited fragments. Sequences from top to bottom: guide RNA targeting SF2 gene, reference SF2 sequence, clones 1-3 showing variable deletions caused by NHEJ repair. (d) Negative geotaxis of F1 *E49>TDP-43, Cas9; U6-gRNA* males over an aging time-course. Y axis shows % flies (average of three trials) crossing a 10cm threshold after 10 second (negative geotaxis testing). X axis indicates time point (7 days or 28 days post eclosion). Bar color indicates gene targeted by gRNA. * P < 0.001 by Student’s T test in comparisons to GFP control. † P < 0.001 by Student’s T test in comparisons to *lst* control.

As a first test of the efficacy and specificity of our cell type specific gRNA targeting strategy, we dissected thoracic ganglia, which contain motor neurons vs gut as a control. DNA from these tissues was extracted and the region flanking the gRNA target sites were queried by PCR and sequencing. Despite the presence of unedited non-motor neuron cells in the thoracic ganglion sample, we were able to detect and sequence internal deletions in the region that is targeted by the *SF2/SRSF1* gRNA. In contrast, such deletions were not detected from gut (**Fig 2B and 2C**).

We next tested the effects on negative geotaxis behavior of the gRNAs against GFP, *lst, mks/importin-7* and *SF2/SRSF1*. We quantified negative geotaxis behavior over the course of a 4 week aging time period and found that relative to the gRNAs targeting GFP or *lst*, the gRNA that targets the *SF2/SRSF1* gene suppressed the effects of TDP-43 in animals that were 7, 21 (not shown) or 28 days old (**Fig 2D**). Similarly, with *msk/importin-7*, each of two independent gRNAs were sufficient to suppress the negative geotaxis defect caused by TDP-43 expression in motor neurons, with *msk* gRNA1 showing significant effects at 7,21 (not shown) and 28 days and gRNA2 showing significant effects at 21 (not shown) and 28 days of age (**Fig 2D**).

### Suppressors prevent age-dependent decline of negative geotaxis behavior

Because neurodegenerative phenotypes exhibit a marked age-dependence, we tested whether our identified suppressors were capable of ameliorating the effects of motor neuron TDP-43 toxicity over an aging time course. For this experiment, we tested 20 of the suppressors that had been equilibrated into a common genetic background (**Fig S1; Table 1**). These were selected to span each of the categories of gene function among the hits from the primary screen. In parallel, we tested a series of 6 negative controls shRNAs. As negative controls, we used UAS-shRNAs against three genes (*ecd, SirT7* and *CG8005*) that had not impacted negative geotaxis in our primary screen. And we also tested 3 different shRNA lines that targeted heterologous transcripts that are not present as endogenous sequences in the fly genome (mCherry, GFP, and Luciferase). Together, these 6 shRNAs controlled for several potential confounds. First, they controlled for the possibility that providing a second UAS-driven transgene might provide functionally relevant reduction in the levels of the *UAS-TDP-43* transgene. We view this possibility as unlikely because with several of our shRNA suppressors, we observe robust expression of GFP in animals that contain the *E49-Gal4* driver, *UAS-GFP* and a *UAS-shRNA* compared with controls that do not contain the shRNA (**Fig S2**). In addition to providing a robust set of controls for the possibility of titrating the expression of *UAS-TDP-43*, these 6 shRNAs also controlled for the possibility that overwhelming the shRNA silencing system might impact TDP-43 neurotoxicity. As with the 20 shRNA suppressors, the 6 control shRNAs are inserted at the common AttP2 PhiC31 landing site, and were equilibrated into the same genetic background.

Negative geotaxis behavior was quantified among F1 males for each of these 26 crosses beginning 1 week after eclosion, and every 7 days thereafter (**Fig 3**). Each of the six negative control shRNAs exhibited severe defects in locomotion at 1 week post eclosion, consistent with their failure to suppress the geotaxis assay in the primary screen. Animals carrying each of these control shRNA also exhibited rapid deterioration in climbing ability at subsequent age time points. By the end of 6 weeks, most of the surviving flies in the negative control groups were largely incapable of climbing (**Fig 3**, red data points). In contrast, 19 out of the 20 shRNA lines that we identified as suppressors in the primary screen exhibited robust climbing ability, with nearly 100% of animals crossing the 10cm distance within 10 seconds. And remarkably, this amelioration of the climbing defects caused by TDP-43 expression in motor neurons was largely resistant to age over a 6 week period (**Fig 3**). The only exception to this is the shRNA targeting the *sbr/NXF1* gene, which was identified as a suppressor in the primary screen (**Table 1; S2 Table**) but whose suppression was eliminated when it was crossed into the w^1118^ (iso CJ1) background (not shown).

**Fig 3.**
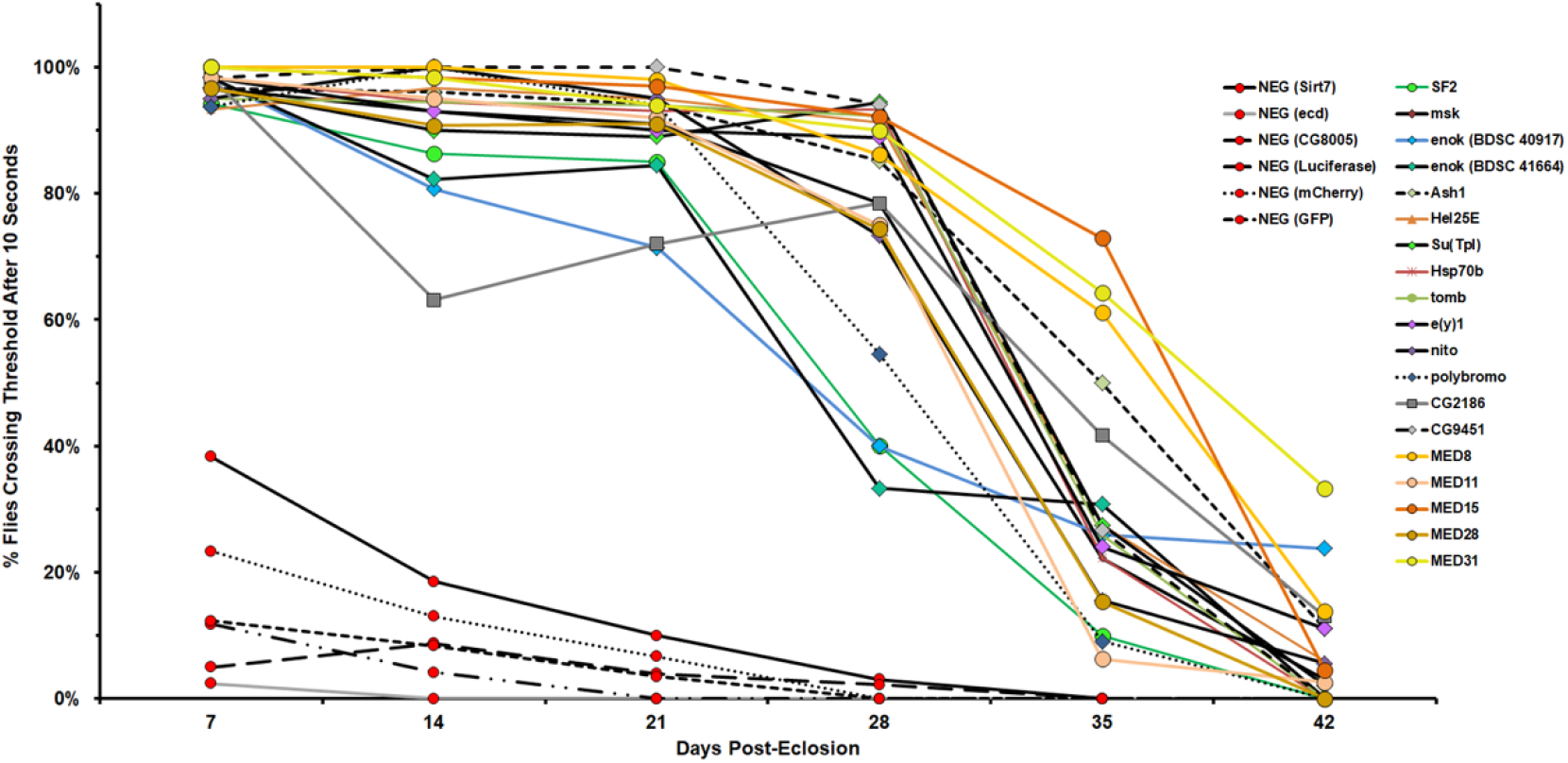
shRNA suppressors rescue the negative geotaxis defect caused by motorneuron-TDP-43 and prevent further age-dependant decline. 25 shRNAs were introgressed into a common wild type *Drosophila* genetic background w^1118^ (iso CJ1), a derivative of CS, and crossed to the TDP-43 screen flies described in Fig. 1a. Negative geotaxis was assayed among F1 progeny of each cross were beginning 7 days after eclosion and each week thereafter. Negative control shRNAs (Red circles) exhibited severe climbing defects that further deteriorated with each subsequent week. Suppressor shRNAs (non-red data points) ameliorated the effects of TDP-43 in young animals (7 day old) and prevented decline in climbing behavior until the 28 day time point or later.

### SF2/SRSF1 and the majority of shRNAs tested strongly suppresses TDP-43 toxicity in a glial context

The primary shRNA screen described above was conducted with TDP-43 expression in motor neurons. But there is a growing recognition of the importance of glial cells in neurodegeneration [37], and we and others have documented that glia play a major role in fly models that involve TDP-43 [38–41]. We therefore wondered whether any of the TDP-43 suppressors that we had identified in neurons also would ameliorate the toxicity of TDP-43 when expressed in glial cells. To test this, we selected a panel of the shRNA suppressors targeting 10 genes that span the functional categories identified. To test the effects of knocking each of these 10 genes in flies that express a non-codon optimized TDP-43 transgene in glia (**see S5 Table**), we made use of an inducible glial expression system in which a temperature sensitive Gal80 repressor is used to modulate the expression of TDP-43 and the shRNA, under control of the Repo-Gal4 that expresses in most glial cells. We have previously shown that inducing pan glial TDP-43 expression after development is sufficient to greatly reduce lifespan, and cause both glial death and non-cell autonomous toxicity of TDP-43 expressing glia that also kills adjacent neurons[41]. We crossed these tub-*GAL80TS, Repo-Gal4; UAS-TDP-43* animals (hereafter referred to as *Repo*^*TS*^ >*TDP43*) with each of the *UAS-shRNAs* and examined lifespans of the F1 offspring after heat-shift induction of TDP-43.

Control flies expressing an shRNA against the ecd gene (shRNA 41676), which did not impact the motor neuron phenotypes with *E49>TDP-43* also had no impact on the severely shortened lifespan of *Repo*^*TS*^ >*TDP-43*. Two independent experiments in which this control shRNA were expressed exhibited median survival of ∼11 days (**Fig. 4** red lines). In contrast, we observe significant lifespan extension with 9/10 shRNAs that target genes identified in the motor neuron suppressor screen. The effect was modest with 2/10 shRNA suppressors (*polybromo, ash1* – **Fig. 4** gray lines, not significant for nito) and was more robust for 6/10 (**Fig. 4**). The most potent suppressor in this glial context was *SF2/SRSF1*, which yielded a median survival of ∼22 and ∼29 days in independently replicated experiments (**Fig. 4** green lines). Despite the expression of TDP-43 in most glia, the lifespan of animals that express *SF2/SRSF1* shRNAs is near that of wild type animals under these temperature conditions (30°C). It also is worth noting that one shRNA, against the *Chd1* gene, was identified as a suppressor in the context of motor-neuron toxicity (**Table 1**) but has the opposite sign in the context of glial TDP-43 toxicity. The shRNA against *Chd1* significantly decreased lifespan of the *Repo*^*TS*^ >*TDP-43* animals. Thus, action of this gene on TDP-43 toxicity may be cell type dependent.

**Fig 4.**
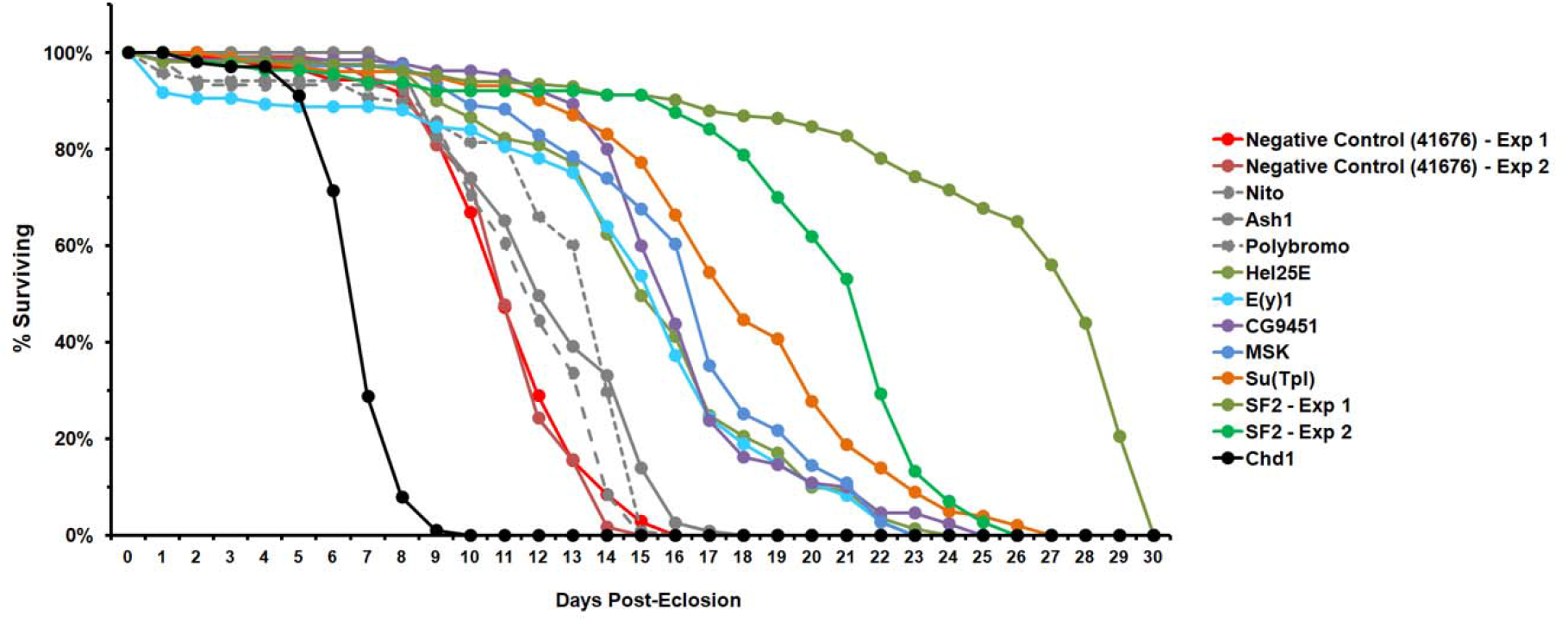
Multiple suppressors, including *SF2/SRSF1*, potently extend lifespan of flies expressing TDP43 in glia. 11 shRNAs that had been equilibrated into a common genetic background w^1118^ (iso CJ1) were crossed to the *RepoTS>TDP43* strain that expresses a UAS-TDP-43 in glia under control of a temperature sensitive Gal80 and the Repo-Gal4 line. These animals induce human TDP-43 in glia after being shifted to 30°C, the non-permissive temperature. F1 males and virgin females were placed into separate vials for aging (20 animals per vial). Time since induction (shift to 30°C) is shown in the X-axis. % survival is shown on the Y-axis. Red data points show 2 independent replicates with a negative control shRNA (shRNA to ecd, introgressed from BDSC 41676) which exhibit nearly complete mortality by 2 weeks. The *Chd1* shRNA (black data points) exacerbates mortality in this glial context, which is in contrast to its effect in motor neurons. *Nito* (gray large dash line) did not extend lifespan significantly after adjusting for multiple testing, nor was the potential effect large. The *ash1* and *polybromo* shRNAs (gray solid and small dashed lines) yield modest but significant increases in lifespan. Each of the other shRNAs (colored data points) increased lifespan dramatically including two independent experiments where the *SF2* shRNA extended lifespan by 2.5-3 fold. Statistical significance (indicate by * on graph) was determined by Kaplan-Meier estimator pairwise comparisons to the negative control, using Bonferroni correction for multiple testing (k = 11, P < 0.004 for significance). P values were <0.00001 for all comparisons except *nito* (P = 0.016).

## Discussion

Overexpression of either wild type or disease alleles of TDP-43 triggers concentration dependent TDP-43 protein aggregation and can lead to TDP-43 clearance from the nucleus and appearance of hyperphosphorylated cytoplasmic inclusions. This approach has been used in yeast [13,14], mammalian cell culture [42–44], and animal models including nematodes [45], flies [25,38,40,41,46,47], and rodents [48–52], and has led to identification of downstream functional defects that have been validated in human tissues. These include defects in splicing [4,6,7,53], nuclear-cytoplasmic localization [17,22,33], DNA damage signaling [40,41,54,55] and expression of retrotransposons and endogenous retroviruses [10,40,41,56–60]. Many of the above findings are derived from high-throughput screens to identify genetic modifiers of TDP-43 toxicity. But to date, no unbiased screen has been conducted in the context of an age-dependent neurological phenotype that emerges from a functioning neural circuit. Moreover, no such screen has been attempted in the context of glia, which play a fundamental role in disease progression [37].

We used over-expression of human TDP-43 in a subset of *Drosophila* motor neurons as the basis for an shRNA knockdown screen to identify suppressors of TDP-43 neurotoxicity. Young (newly eclosed) adult flies with such motor neuron expression of TDP-43 exhibit mild defects in climbing behavior in a negative geotaxis assay, but this defect becomes dramatically worse with age. Using this approach, we screened a collection of shRNAs that covered approximately 15-20% of protein coding genes in the *Drosophila* genome, and found that approximately 1% of shRNAs can abrogate TDP-43 toxicity. For the majority of these, shRNA knockdown also potently prevented the age-dependent decline in locomotion seen in controls. Additionally, for most of the cases tested, effects on lifespan from TDP-43 expression in glia also was ameliorated.

The majority of suppressors described here have not previously been implicated in neurodegeneration. But several of the hits have been identified in previous screens for TDP-43 suppressors, and a number of others play roles in cellular pathways that have established impact on TDP-43 related neurodegeneration. For example, we identified 8 genes with roles in chromatin remodeling. These include several specific genes that have been found in previous screens. *Chd1* and *moira*, for instance, are chromatin remodeling factors that were identified [23] in a previous screen in flies for modifiers of a TDP-43 induced rough eye phenotype. It is worth noting, however, that in this previous study, downregulation of *Chd1* exacerbated the toxicity of TDP-43 to photoreceptor neurons, leading to a more severe rough eye phenotype. This is the opposite sign of effect that we observed in the context of our motor-neuron based behavioral screen. On the other hand, when we screened hits in the context of effects of glial TDP-43 on lifespan, we found that *Chd1* knockdown exacerbated the toxicity, consistent with the sign of effect seen in the previous study. This drives home the importance of cellular context, and the importance of testing impact within both glia and neurons.

In the case of the 8 chromatin modulators identified as suppressors, their functional roles may also provide some mechanistic insight. With the exception of *Chd1*, which promotes chromatin silencing, the 7 other genes in this category all promote open chromatin. *E(y)3, polybromo, ash1, moira, br* and *br140* are all part of the *trithorax* and SWI/SNF (Brahma) complexes and *enok* is a histone acetyltransferase that interacts with *trithorax* group genes and also promotes open chromatin. With each of these, TDP-43 toxicity is ameliorated by shRNA knock down, suggesting that chromatin silencing normally helps keep TDP-43 toxicity in check. This idea is consistent with the hypothesis that loss of silencing of retrotransposons and endogenous retroviruses contributes to TDP-43 toxicity [10,40,41,56,57,60].

A second category of suppressor identified here are factors involved in basal transcription. This includes 8 components of the Mediator complex, and several basal transcription factors. Although it is difficult to distinguish specific vs artifactual causes for identification of factors regulating fundamental aspects of transcription, it is worth pointing out that we also identified the transcription elongation factor, *Su(Tpl)*, which was found in a previous rough eye suppressor screen [61]. In that study, the authors present compelling evidence that in the presence of TDP-43 pathology, elongator complexes aberrantly express small nucleolar RNAs. Our identification of *Su(Tpl)* suggests that a similar process could be at play in motor neurons and glia.

The third major category of suppressor includes genes with known roles in nuclear import/export control, a pathway that was also convergently identified in several previous screens [17,22], the idea that defects in nuclear-cytoplasmic shuttling may be responsible for cytoplasmic mis-localization of TDP-43. We were intrigued that *msk*, one of the nuclear import factors that we identified is the *Drosophila* orthologue of Importin 7, a nuclear import protein that has a role in nuclear entry of HIV retrovirus [62,63] and LINE1 retrotransposons [64]. Thus, as with the chromatin silencers discussed above, this provides a possible link to involvement of retrotransposons and endogenous retroviruses.

Finally, we identified *SF2/SRSF1* as a particularly robust suppressor. We found that knock down of *SF2/SRSF1* abrogated the locomotion defect caused by TDP-43 expression in motor neurons, prevented the age-dependent decline in climbing ability out to 6-weeks of age, and potently extended lifespan in animals that over-express TDP-43 in glial cells. In fact, glial knock down of *SF2/SRSF1* is sufficient to all but eliminate the severe lifespan effect seen in animals that overexpress TDP-43 in glia. This RNA binding protein functions in both splicing and as a nuclear export of adaptor for RNA export. It has previously been identified in a *Drosophila* screen for suppressors of a rough eye phenotype caused by *C9orf72* hexanucleotide expansion repeat RNA [24]. In that study, the authors determined that *SF2/SRSF1* knock down could suppress the toxicity of a *C9orf72* repeat RNA that encoded dipeptide repeats. This was demonstrated both in the Drosophila context and in a co-culture system in which astrocytes from *C9orf72* derived patient fibroblasts are grown with motor neurons. In the induced astrocyte context, knock down of *SF2/SRSF1* was sufficient to ameliorate the toxicity of patient derived astrocytes to motor neurons. This is reminiscent of our observation of highly potent suppression of the effect on lifespan of glial TDP-43 expression in flies, an effect that we previously have documented to involve toxicity of the glia to neurons [41]. In the case of *C9orf72*, the effect of *SF2/SRSF1* knock down was shown to be mediated by modulation of the export of the repeat RNA to the cytoplasm. The reduced repeat RNA export in turn was proposed to reduce the repeat associated translation to produce toxic dipeptide repeats. In fact, *SF2/SRSF1* knockdown was not sufficient to significantly ameliorate the toxicity of arginine rich dipeptides expressed from non-repeat RNAs[24], consistent with the interpretation that the relevant impact of *SF2/SRSF1* is to aid export of the repeat RNA to the cytoplasm

Our identification of *SF2/SRSF1* as a suppressor of TDP-43 toxicity in neurons and glial cells in *Drosophila* suggests convergence with the effects previously observed on the *C9orf72* repeat. In principle, this convergence could reflect the fact that *C9orf72* patients typically exhibit TDP-43 cytoplasmic inclusions [13]. Indeed in patient brain tissues, TDP-43 protein pathology often co-localizes with that of the GR dipeptide repeat [65]. Furthermore, animal models of *C9orf72* expansion toxicity also can exhibit TDP-43 proteinopathy [66]. On the other hand, we find that knock down of *SF2/SRSF1* has strong suppressive effects on the toxicity of TDP-43 that occurs in the absence of any *C9orf72* repeat RNAs. So in this case, effects of *SF2/SRSF1* knock down on toxicity of TDP-43 is independent of any impact on *C9orf72* repeat RNA export.

We have used a genetic screening approach to identify suppressors of TDP-43 toxicity in Drosophila. Unlike previous screens, we used an age-dependent behavioral defect as a primary readout. This age-dependent effect on neurobehavioral readout may enrich for effects that are related to the age-dependent degenerative rather than neurodevelopmental effects, but also is sensitive to disruptions to the interactions among each of the relevant cell types to a locomotion circuit, including motor neurons and glia. Our secondary screen in for suppression of toxicity to glia also reveals that many of the same genetic pathways participate in toxicity to glia and neurons. This approach shows convergence with several key findings from previous screens, namely involvement of chromatin silencing and nuclear/cytoplasmic shuttling, but also identifies genes not previously associated with neurodegeneration. Finally, the identification of *SF2/SRSF1* provides a point of convergence between TDP-43 proteinopathy and *C9orf72* familial ALS/FTD and points to unexplored mechanistic underpinnings of that overlap.

## Materials and Methods

### Negative Geotaxis of shRNA and CRISPR/Cas9 Screening Systems

Male *GAL80;E49>TDP-43* animals were crossed to females that contained either the UAS-shRNA or the gRNA and UAS-Cas9. In each case, Male F1 animals that had lost the Gal80 (both contained the E49-Gal4, the UAS-TDP-43 and either the UAS-shRNA or the gRNA and the UAS-Cas9) were collected shortly after eclosing and transferred to fresh vials with food. Behavioral testing took place between 12PM and 4PM in an isolated environmentally controlled room. Lighting conditions were kept as consistent as possible for all geotaxis screening. For the assay, a maximum of 20 flies were quickly transferred (without being anesthetized) by tapping the housing vial and transferring to an assay vial without food. Flies were placed into two 95mm vials whose open mouths were held together by black plastic gaskets. A frame capable of holding ten sets of vial pairs was used to simultaneously tap flies to the bottom of the vials. A timer and video camera were started prior to tapping the flies down. Video of climbing activity was recorded for three trials per experiment. Scoring of geotaxis was performed by determining the percentage of the flies in each vial that crossed to the top vial (10cm) within 10 seconds (recording proceeded for 15 to 20 seconds).

### Lifespan in Glial Expression System

Male and virgin female flies were collected over a period of 72 hours. Groups of approximately 20 flies were placed in each of 3 separate vials (n=60 for most assayed genotypes). Males and females were tested separately. Flies were kept in temperature and humidity controlled incubators. To induce TDP-43 expression in glia, the flies were transferred to a 30°C incubator. Flies were flipped to vials of new food every other day to avoid mortality from becoming stuck in food and dead flies were removed and counted every day.

### CRISPR Genome Editing

Candidate gRNAs were designed by a publicly available computational prediction method https://www.flyrnai.org/crispr3/web/ [67]. For each gene of interest, 5 candidate sequences were chosen from the program output, and further filtered. A positive strand target and a negative strand target were chosen. Sequences with internal U6 terminators were excluded. All sequences listed include PAM sites (3 terminal base pairs) and were excluded from ordered oligos. Sequences were chosen on the basis of high likelihood of cutting (Housden and Machine scores) and having a high probability of inducing a frameshift after NHEJ repair. Sequences used can be found in **S3 Table**.

Oligos (**S4 Table**) were ordered from IDTDNA and included overhangs for Bbs1 restriction sites. Oligos were annealed and cloned into a linearized pCFD3.1 vector [68], cloned and purified using Qiagen EndoFree plasmid Maxiprep kit, sequenced for confirmation and sent for microinjection to BestGene Drosophila Injection Services. Transgenic U6-gRNA flies were introgressed and homogenized (pCFD3.1 was integrated into AttP2 site on Chromosome 3) by screening for the *white* phenotype rescue. CRISPR cutting in motor neurons was assessed as described in **Fig 4B**. Screening primers can be found in **S3 Table**.

### Cas9 Deletion Confirmation PCR, Fragment Cloning and Sequencing

Cas9 gRNA F1 Male flies were anesthetized and pinned to a dissecting plate. The gut and thoracic ganglia were separately dissected from 10 flies and placed into DNA extraction buffer, homogenized, and treated with Proteinase K. PCR amplification using KAPA2G Fast HotStart Genotyping Mix (Kapa Biosystems) with flanking oligos (**S4 Table**) was performed for 30 cycles then amplification fragments were evaluated by agarose gel electrophoresis (1.8% w/v TE buffer). Bands indicated in **Fig. 2B** were excised with a scalpel, purified using Zymoclean Gel DNA Recovery Kit (Zymo Research) and cloned into TOPO® TA Cloning vectors (Thermo Fisher Scientific).

### Confocal Imaging of E49>GFP, shRNA Thoracic Ganglia

F1 males flies from E49>GFP, UAS-shRNA crosses were anesthetized, then briefly dipped in 95% EtOH and pinned to a dissecting plate. The thoracic ganglia from at least 3 flies was removed and immediately placed in cold 4% PFA/1x PBST (phosphate buffered saline + 2% Triton X-100 + 10% Normal Goat Serum) for 15 minutes, then degassed in fixative at room temperature for 45 minutes. 3 washes with PBST (10 minutes each) were performed, and ganglia were incubated with a mouse anti-GFP 1° antibody (1:250 dilution MilliporeSigma #11814460001) in 1x PBST (0.2% Triton X-100 + 10% Normal Goat Serum) overnight at 4°C. Ganglia were then washed 3 times (10 minutes each) with 1x PBS and then incubated with 2° antibody (1:250 dilution of Alexa Fluor 488-conjugated goat anti-mouse, Thermo Fisher Scientific #A-11029) in 1x PBST (0.2% Triton X-100 + 10% Normal Goat Serum) for 1 hour at room temperature, then washed with 1x PBST 3 times (10 minutes each) and placed on a concave dissecting dish. The PBST was removed, and the tissue was briefly rinsed with 1x PBS, then as much of the liquid was removed as possible with a pipette before adding FocusClear™(CelExplorer) directly to the tissue. After tissue clearing occurred, the ganglia were transferred to a slide with a drop of ProLong™Diamond AntiFade Mountant with DAPI (Thermo Fisher Scientific) and a glass coverslip placed over the preparation. The cover was glass was sealed with nail polish and allowed to set overnight in a dark 4°C refrigerator. Images were generated with a Zeiss Confocal microscope.

### Fly Stocks

See **S5 Table** for all *Drosophila* stocks used in this study. TRiP stocks were not treated for the possible presence of *Wolbachia*.

## Acknowledgements and Author Contributions

The authors would like to thank Rachel Cuozzo and Chandana Kochath for their help with screening and stock maintenance. We would also like to thank Yung-Heng Chang, Meng-Fu Shih, Sarah Krupp, Richard Keegan, Lillian Talbot,Roger B. Sher and Maurice Kernan for their critique, invaluable advice and assistance. Caitlin Maher Dubnau provided assistance building the apparatus for parallel testing of multiple strains for negative geotaxis.

J.A. wrote manuscript, designed experiments, and carried out screening, genetics, and analyzed data. E.E. carried out screening and analyzed data. M.T. created constructs and helped with screening. J.D. designed experiments, analyzed data, and edited the manuscript.

This work was supported by grants to J.D. from NINDS (R01NS091748), the NIA (RF1AG057338), and the ALS Ride for Life charitable foundation

## Competing Interests

The authors declare no competing interests.

## Supplemental Material

**S1 Table** – Retesting of non-rescuing lines to determine false positive rate.

**S2 Table** – Comprehensive screen data. Sheet 1: first screen of full TRiP line assembly. Sheet 2: retest summary of candidate rescues for validation.

**Fig S1.**
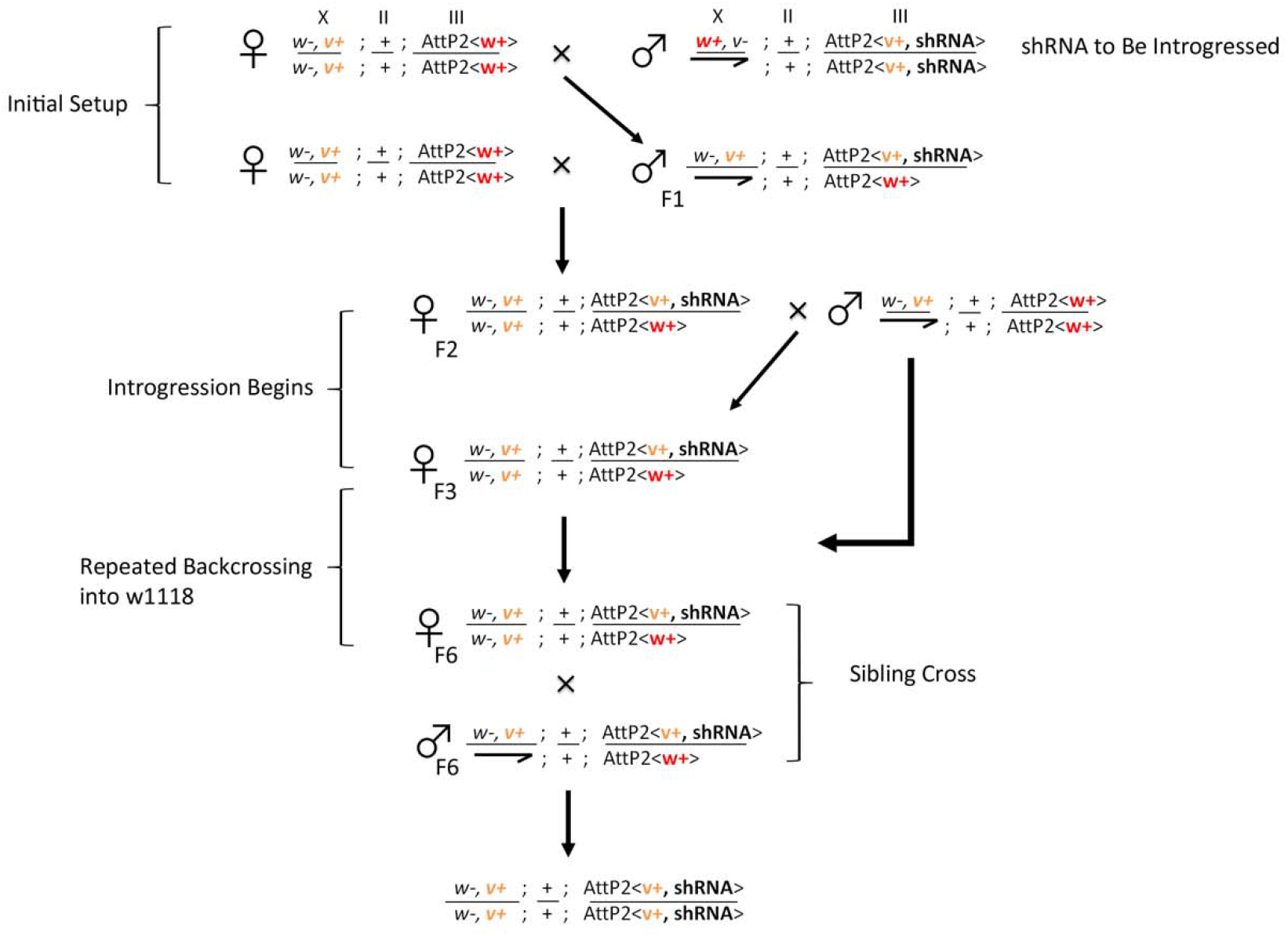
Introgression scheme. TRiP lines carry the vermillion (v) marker on their AttP2 integration site. Initial setup: virgin w^1118^ with a miniwhite transgene inserted into AttP2 were crossed to males from the desired TRiP line. F1 offspring males heterozygous on the third chromosome were again crossed to virgins from the w^1118^ background to yield F2 virgin females homozygous on the X for w- and heterozygous on the 3^rd^ chromosome for introgression. Introgression and backcrossing: F2 virgin females heterozygous on the third (very light orange eyes) were backcrossed to males from the w1118 background. This was repeated with the every subsequent generation (F3-F6) of females heterozygous on the third to gradually replace more of the 3^rd^ chromosome containing the AttP2<shRNA> with the third chromosome from the w1118 background. Sibling cross: after 6 generations of introgression, male and female siblings with light orange eyes were crossed to each other, and F7 offspring with white eyes were selected (completely lacking miniwhite in the AttP2 site, indicating presence of shRNA).

**Fig S2.**
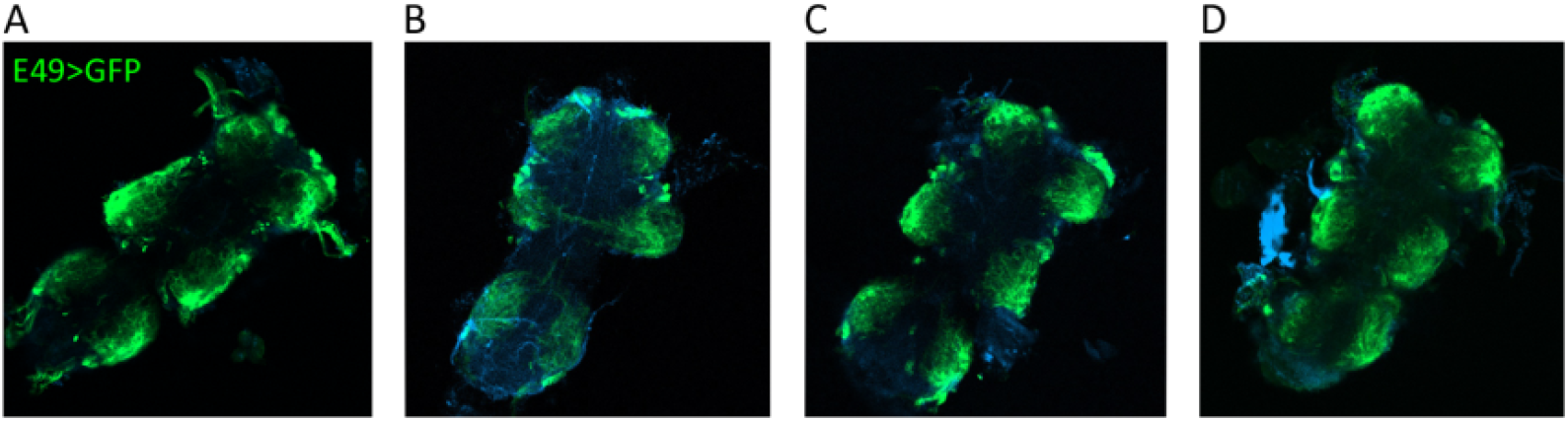
Confocal microscopy of thoracic ganglia from E49>GFP flies expressing rescuing shRNAs. All images are single-plane on 10X objective. shRNAs were (A) *msk*, (B) *Su(Tpl)*, (C) *SF2*, (D) *Polybromo*.

**S3 Table** – Sheet 1: computational output showing gRNA properties. Sheet 2: gRNA oligonucleotide sequences ordered for cloning into pCFD3.1 vector.

**S4 Table** – PCR Primers for Cas9 cutting validation.

**S5 Table** – *Drosophila* stocks used in this study.

## Notes

### Competing Interest Statement

The authors have declared no competing interest.

## References

1. Al-Chalabi A, Hardiman O. The epidemiology of ALS: a conspiracy of genes, environment and time. Nat Rev Neurol. 2013;9: 617–628. doi:10.1038/nrneurol.2013.203

2. Hardiman O, Al-Chalabi A, Chio A, Corr EM, Logroscino G, Robberecht W, et al. Amyotrophic lateral sclerosis. Nat Rev Dis Primers. 2017;3: 17071. doi:10.1038/nrdp.2017.71

3. Ou SH, Wu F, Harrich D, García-Martínez LF, Gaynor RB. Cloning and characterization of a novel cellular protein, TDP-43, that binds to human immunodeficiency virus type 1 TAR DNA sequence motifs. J Virol. 1995;69: 3584–3596. doi:10.1128/JVI.69.6.3584-3596.1995

4. Buratti E, Baralle FE. Multiple roles of TDP-43 in gene expression, splicing regulation, and human disease. Front Biosci. 2008;13: 867–878. doi:10.2741/2727

5. Buratti E, Baralle FE. The multiple roles of TDP-43 in pre-mRNA processing and gene expression regulation. RNA Biology. 2010;7: 420–429. doi:10.4161/rna.7.4.12205

6. Tollervey JR, Curk T, Rogelj B, Briese M, Cereda M, Kayikci M, et al. Characterizing the RNA targets and position-dependent splicing regulation by TDP-43. Nat Neurosci. 2011;14: 452–458. doi:10.1038/nn.2778

7. Lukavsky PJ, Daujotyte D, Tollervey JR, Ule J, Stuani C, Buratti E, et al. Molecular basis of UG-rich RNA recognition by the human splicing factor TDP-43. Nat Struct Mol Biol. 2013;20: 1443–1449. doi:10.1038/nsmb.2698

8. Ling JP, Pletnikova O, Troncoso JC, Wong PC. TDP-43 repression of nonconserved cryptic exons is compromised in ALS-FTD. Science. 2015;349: 650–655. doi:10.1126/science.aab0983

9. Saldi TK, Ash PE, Wilson G, Gonzales P, Garrido-Lecca A, Roberts CM, et al. TDP-1, the Caenorhabditis elegans ortholog of TDP-43, limits the accumulation of double-stranded RNA. EMBO J. 2014;33: 2947–2966. doi:10.15252/embj.201488740

10. Liu EY, Russ J, Cali CP, Phan JM, Amlie-Wolf A, Lee EB. Loss of Nuclear TDP-43 Is Associated with Decondensation of LINE Retrotransposons. Cell Reports. 2019;27: 1409-1421.e6. doi:10.1016/j.celrep.2019.04.003

11. Yokoseki A, Shiga A, Tan C-F, Tagawa A, Kaneko H, Koyama A, et al. TDP-43 mutation in familial amyotrophic lateral sclerosis. Ann Neurol. 2008;63: 538–542. doi:10.1002/ana.21392

12. Sreedharan J, Blair IP, Tripathi VB, Hu X, Vance C, Rogelj B, et al. TDP-43 mutations in familial and sporadic amyotrophic lateral sclerosis. Science. 2008;319: 1668–1672. doi:10.1126/science.1154584

13. Byrne S, Elamin M, Bede P, Shatunov A, Walsh C, Corr B, et al. Cognitive and clinical characteristics of patients with amyotrophic lateral sclerosis carrying a C9orf72 repeat expansion: a population-based cohort study. The Lancet Neurology. 2012;11: 232–240. doi:10.1016/S1474-4422(12)70014-5

14. Johnson BS, McCaffery JM, Lindquist S, Gitler AD. A yeast TDP-43 proteinopathy model: Exploring the molecular determinants of TDP-43 aggregation and cellular toxicity. Proceedings of the National Academy of Sciences. 2008;105: 6439–6444. doi:10.1073/pnas.0802082105

15. Couthouis J, Hart MP, Shorter J, DeJesus-Hernandez M, Erion R, Oristano R, et al. A yeast functional screen predicts new candidate ALS disease genes. Proceedings of the National Academy of Sciences. 2011;108: 20881–20890. doi:10.1073/pnas.1109434108

16. Sun Z, Diaz Z, Fang X, Hart MP, Chesi A, Shorter J, et al. Molecular Determinants and Genetic Modifiers of Aggregation and Toxicity for the ALS Disease Protein FUS/TLS. Weissman JS, editor. PLoS Biol. 2011;9: e1000614. doi:10.1371/journal.pbio.1000614

17. Jovičić A, Mertens J, Boeynaems S, Bogaert E, Chai N, Yamada SB, et al. Modifiers of C9orf72 dipeptide repeat toxicity connect nucleocytoplasmic transport defects to FTD/ALS. Nat Neurosci. 2015;18: 1226–1229. doi:10.1038/nn.4085

18. Elden AC, Kim H-J, Hart MP, Chen-Plotkin AS, Johnson BS, Fang X, et al. Ataxin-2 intermediate-length polyglutamine expansions are associated with increased risk for ALS. Nature. 2010;466: 1069–1075. doi:10.1038/nature09320

19. Van Damme P, Veldink JH, van Blitterswijk M, Corveleyn A, van Vught PWJ, Thijs V, et al. Expanded ATXN2 CAG repeat size in ALS identifies genetic overlap between ALS and SCA2. Neurology. 2011;76: 2066–2072. doi:10.1212/WNL.0b013e31821f445b

20. Liu X, Lu M, Tang L, Zhang N, Chui D, Fan D. ATXN2 CAG repeat expansions increase the risk for Chinese patients with amyotrophic lateral sclerosis. Neurobiology of Aging. 2013;34: 2236.e5-2236.e8. doi:10.1016/j.neurobiolaging.2013.04.009

21. Boeynaems S, Bogaert E, Michiels E, Gijselinck I, Sieben A, Jovičić A, et al. Drosophila screen connects nuclear transport genes to DPR pathology in c9ALS/FTD. Sci Rep. 2016;6: 20877. doi:10.1038/srep20877

22. Freibaum BD, Lu Y, Lopez-Gonzalez R, Kim NC, Almeida S, Lee K-H, et al. GGGGCC repeat expansion in C9orf72 compromises nucleocytoplasmic transport. Nature. 2015;525: 129–133. doi:10.1038/nature14974

23. Berson A, Sartoris A, Nativio R, Van Deerlin V, Toledo JB, Porta S, et al. TDP-43 Promotes Neurodegeneration by Impairing Chromatin Remodeling. Current Biology. 2017;27: 3579-3590.e6. doi:10.1016/j.cub.2017.10.024

24. Hautbergue GM, Castelli LM, Ferraiuolo L, Sanchez-Martinez A, Cooper-Knock J, Higginbottom A, et al. SRSF1-dependent nuclear export inhibition of C9ORF72 repeat transcripts prevents neurodegeneration and associated motor deficits. Nat Commun. 2017;8: 16063. doi:10.1038/ncomms16063

25. Yusuff T, Chatterjee S, Chang Y-C, Sang T-K, Jackson GR. Codon-optimized TDP-43-mediated neurodegeneration in a Drosophila model for ALS/FTLD. Neuroscience; 2019 Jul. doi:10.1101/696963

26. Gordon MD, Scott K. Motor Control in a Drosophila Taste Circuit. Neuron. 2009;61: 373–384. doi:10.1016/j.neuron.2008.12.033

27. Azpurua J, Mahoney RE, Eaton BA. Transcriptomics of aged Drosophila motor neurons reveals a matrix metalloproteinase that impairs motor function. Aging Cell. 2018;17. doi:10.1111/acel.12729

28. Ni J-Q, Zhou R, Czech B, Liu L-P, Holderbaum L, Yang-Zhou D, et al. A genome-scale shRNA resource for transgenic RNAi in Drosophila. Nat Methods. 2011;8: 405–407. doi:10.1038/nmeth.1592

29. Gargano J, Martin I, Bhandari P, Grotewiel M. Rapid iterative negative geotaxis (RING): a new method for assessing age-related locomotor decline in. Experimental Gerontology. 2005;40: 386–395. doi:10.1016/j.exger.2005.02.005

30. Allen BL, Taatjes DJ. The Mediator complex: a central integrator of transcription. Nat Rev Mol Cell Biol. 2015;16: 155–166. doi:10.1038/nrm3951

31. Berson A, Nativio R, Berger SL, Bonini NM. Epigenetic Regulation in Neurodegenerative Diseases. Trends in Neurosciences. 2018;41: 587–598. doi:10.1016/j.tins.2018.05.005

32. Tibshirani M, Zhao B, Gentil BJ, Minotti S, Marques C, Keith J, et al. Dysregulation of chromatin remodelling complexes in amyotrophic lateral sclerosis. Human Molecular Genetics. 2017;26: 4142–4152. doi:10.1093/hmg/ddx301

33. Kim HJ, Taylor JP. Lost in Transportation: Nucleocytoplasmic Transport Defects in ALS and Other Neurodegenerative Diseases. Neuron. 2017;96: 285–297. doi:10.1016/j.neuron.2017.07.029

34. Archbold HC, Jackson KL, Arora A, Weskamp K, Tank EM-H, Li X, et al. TDP43 nuclear export and neurodegeneration in models of amyotrophic lateral sclerosis and frontotemporal dementia. Sci Rep. 2018;8: 4606. doi:10.1038/s41598-018-22858-w

35. Boehringer A, Garcia-Mansfield K, Singh G, Bakkar N, Pirrotte P, Bowser R. ALS Associated Mutations in Matrin 3 Alter Protein-Protein Interactions and Impede mRNA Nuclear Export. Sci Rep. 2017;7: 14529. doi:10.1038/s41598-017-14924-6

36. Cooper-Knock J, Robins H, Niedermoser I, Wyles M, Heath PR, Higginbottom A, et al. Targeted Genetic Screen in Amyotrophic Lateral Sclerosis Reveals Novel Genetic Variants with Synergistic Effect on Clinical Phenotype. Front Mol Neurosci. 2017;10: 370. doi:10.3389/fnmol.2017.00370

37. Philips T, Rothstein JD. Glial cells in amyotrophic lateral sclerosis. Experimental Neurology. 2014;262: 111–120. doi:10.1016/j.expneurol.2014.05.015

38. Estes PS, Daniel SG, Mccallum AP, Boehringer AV, Sukhina AS, Zwick RA, et al. Motor neurons and glia exhibit specific individualized responses to TDP-43 expression in a Drosophila model of amyotrophic lateral sclerosis. Disease Models & Mechanisms. 2013;6: 721–733. doi:10.1242/dmm.010710

39. Diaper DC, Adachi Y, Lazarou L, Greenstein M, Simoes FA, Di Domenico A, et al. Drosophila TDP-43 dysfunction in glia and muscle cells cause cytological and behavioural phenotypes that characterize ALS and FTLD. Human Molecular Genetics. 2013;22: 3883–3893. doi:10.1093/hmg/ddt243

40. Krug L, Chatterjee N, Borges-Monroy R, Hearn S, Liao W-W, Morrill K, et al. Retrotransposon activation contributes to neurodegeneration in a Drosophila TDP-43 model of ALS. Feschotte C, editor. PLoS Genet. 2017;13: e1006635. doi:10.1371/journal.pgen.1006635

41. Chang Y-H, Dubnau J. The Gypsy Endogenous Retrovirus Drives Non-Cell-Autonomous Propagation in a Drosophila TDP-43 Model of Neurodegeneration. Curr Biol. 2019;29: 3135-3152.e4. doi:10.1016/j.cub.2019.07.071

42. Shiina Y, Arima K, Tabunoki H, Satoh J. TDP-43 Dimerizes in Human Cells in Culture. Cell Mol Neurobiol. 2010;30: 641–652. doi:10.1007/s10571-009-9489-9

43. Bilican B, Serio A, Barmada SJ, Nishimura AL, Sullivan GJ, Carrasco M, et al. Mutant induced pluripotent stem cell lines recapitulate aspects of TDP-43 proteinopathies and reveal cell-specific vulnerability. Proceedings of the National Academy of Sciences. 2012;109: 5803–5808. doi:10.1073/pnas.1202922109

44. Liu-Yesucevitz L, Bilgutay A, Zhang Y-J, Vanderwyde T, Citro A, Mehta T, et al. Tar DNA Binding Protein-43 (TDP-43) Associates with Stress Granules: Analysis of Cultured Cells and Pathological Brain Tissue. Bush AI, editor. PLoS ONE. 2010;5: e13250. doi:10.1371/journal.pone.0013250

45. Ash PEA, Zhang Y-J, Roberts CM, Saldi T, Hutter H, Buratti E, et al. Neurotoxic effects of TDP-43 overexpression in C. elegans. Human Molecular Genetics. 2010;19: 3206–3218. doi:10.1093/hmg/ddq230

46. Li Y, Ray P, Rao EJ, Shi C, Guo W, Chen X, et al. A Drosophila model for TDP-43 proteinopathy. Proceedings of the National Academy of Sciences. 2010;107: 3169–3174. doi:10.1073/pnas.0913602107

47. Miguel L, Frébourg T, Campion D, Lecourtois M. Both cytoplasmic and nuclear accumulations of the protein are neurotoxic in Drosophila models of TDP-43 proteinopathies. Neurobiology of Disease. 2011;41: 398–406. doi:10.1016/j.nbd.2010.10.007

48. Stallings NR, Puttaparthi K, Luther CM, Burns DK, Elliott JL. Progressive motor weakness in transgenic mice expressing human TDP-43. Neurobiology of Disease. 2010;40: 404–414. doi:10.1016/j.nbd.2010.06.017

49. Wils H, Kleinberger G, Janssens J, Pereson S, Joris G, Cuijt I, et al. TDP-43 transgenic mice develop spastic paralysis and neuronal inclusions characteristic of ALS and frontotemporal lobar degeneration. Proc Natl Acad Sci USA. 2010;107: 3858–3863. doi:10.1073/pnas.0912417107

50. Xu Y-F, Gendron TF, Zhang Y-J, Lin W-L, D’Alton S, Sheng H, et al. Wild-Type Human TDP-43 Expression Causes TDP-43 Phosphorylation, Mitochondrial Aggregation, Motor Deficits, and Early Mortality in Transgenic Mice. Journal of Neuroscience. 2010;30: 10851–10859. doi:10.1523/JNEUROSCI.1630-10.2010

51. Xu Y-F, Zhang Y-J, Lin W-L, Cao X, Stetler C, Dickson DW, et al. Expression of mutant TDP-43 induces neuronal dysfunction in transgenic mice. Mol Neurodegeneration. 2011;6: 73. doi:10.1186/1750-1326-6-73

52. Janssens J, Wils H, Kleinberger G, Joris G, Cuijt I, Ceuterick-de Groote C, et al. Overexpression of ALS-Associated p.M337V Human TDP-43 in Mice Worsens Disease Features Compared to Wild-type Human TDP-43 Mice. Mol Neurobiol. 2013;48: 22–35. doi:10.1007/s12035-013-8427-5

53. Arnold ES, Ling S-C, Huelga SC, Lagier-Tourenne C, Polymenidou M, Ditsworth D, et al. ALS-linked TDP-43 mutations produce aberrant RNA splicing and adult-onset motor neuron disease without aggregation or loss of nuclear TDP-43. Proceedings of the National Academy of Sciences. 2013;110: E736–E745. doi:10.1073/pnas.1222809110

54. Guerrero EN, Mitra J, Wang H, Rangaswamy S, Hegde PM, Basu P, et al. Amyotrophic lateral sclerosis-associated TDP-43 mutation Q331K prevents nuclear translocation of XRCC4-DNA ligase 4 complex and is linked to genome damage-mediated neuronal apoptosis. Human Molecular Genetics. 2019;28: 2459–2476. doi:10.1093/hmg/ddz062

55. Mitra J, Guerrero EN, Hegde PM, Liachko NF, Wang H, Vasquez V, et al. Motor neuron disease-associated loss of nuclear TDP-43 is linked to DNA double-strand break repair defects. Proc Natl Acad Sci USA. 2019;116: 4696–4705. doi:10.1073/pnas.1818415116

56. Li W, Jin Y, Prazak L, Hammell M, Dubnau J. Transposable Elements in TDP-43-Mediated Neurodegenerative Disorders. Iijima KM, editor. PLoS ONE. 2012;7: e44099. doi:10.1371/journal.pone.0044099

57. Douville R, Liu J, Rothstein J, Nath A. Identification of active loci of a human endogenous retrovirus in neurons of patients with amyotrophic lateral sclerosis. Ann Neurol. 2011;69: 141–151. doi:10.1002/ana.22149

58. Tam OH, Rozhkov NV, Shaw R, Kim D, Hubbard I, Fennessey S, et al. Postmortem Cortex Samples Identify Distinct Molecular Subtypes of ALS: Retrotransposon Activation, Oxidative Stress, and Activated Glia. Cell Rep. 2019;29: 1164-1177.e5. doi:10.1016/j.celrep.2019.09.066

59. Prudencio M, Gonzales PK, Cook CN, Gendron TF, Daughrity LM, Song Y, et al. Repetitive element transcripts are elevated in the brain of C9orf72 ALS/FTLD patients. Human Molecular Genetics. 2017;26: 3421–3431. doi:10.1093/hmg/ddx233

60. Romano G, Klima R, Feiguin F. TDP-43 prevents retrotransposon activation in the Drosophila motor system through regulation of Dicer-2 activity. BMC Biol. 2020;18: 82. doi:10.1186/s12915-020-00816-1

61. Chung C-Y, Berson A, Kennerdell JR, Sartoris A, Unger T, Porta S, et al. Aberrant activation of non-coding RNA targets of transcriptional elongation complexes contributes to TDP-43 toxicity. Nat Commun. 2018;9: 4406. doi:10.1038/s41467-018-06543-0

62. Zaitseva L, Cherepanov P, Leyens L, Wilson SJ, Rasaiyaah J, Fassati A. HIV-1 exploits importin 7 to maximize nuclear import of its DNA genome. Retrovirology. 2009;6: 11. doi:10.1186/1742-4690-6-11

63. Fassati A. Nuclear import of HIV-1 intracellular reverse transcription complexes is mediated by importin 7. The EMBO Journal. 2003;22: 3675–3685. doi:10.1093/emboj/cdg357

64. Taylor MS, Altukhov I, Molloy KR, Mita P, Jiang H, Adney EM, et al. Dissection of affinity captured LINE-1 macromolecular complexes. eLife. 2018;7: e30094. doi:10.7554/eLife.30094

65. Saberi S, Stauffer JE, Jiang J, Garcia SD, Taylor AE, Schulte D, et al. Sense-encoded poly-GR dipeptide repeat proteins correlate to neurodegeneration and uniquely co-localize with TDP-43 in dendrites of repeat-expanded C9orf72 amyotrophic lateral sclerosis. Acta Neuropathol. 2018;135: 459–474. doi:10.1007/s00401-017-1793-8

66. Chew J, Gendron TF, Prudencio M, Sasaguri H, Zhang Y-J, Castanedes-Casey M, et al. C9ORF72 repeat expansions in mice cause TDP-43 pathology, neuronal loss, and behavioral deficits. Science. 2015;348: 1151–1154. doi:10.1126/science.aaa9344

67. Housden BE, Valvezan AJ, Kelley C, Sopko R, Hu Y, Roesel C, et al. Identification of potential drug targets for tuberous sclerosis complex by synthetic screens combining CRISPR-based knockouts with RNAi. Sci Signal. 2015;8: rs9. doi:10.1126/scisignal.aab3729

68. Port F, Chen H-M, Lee T, Bullock SL. Optimized CRISPR/Cas tools for efficient germline and somatic genome engineering in Drosophila. Proceedings of the National Academy of Sciences. 2014;111: E2967–E2976. doi:10.1073/pnas.1405500111

